# The Endless Frontier? The Recent Upsurge of R&D Productivity in Pharmaceuticals

**DOI:** 10.1101/670471

**Authors:** Fabio Pammolli, Lorenzo Righetto, Sergio Abrignani, Luca Pani, Pier Giuseppe Pelicci, Emanuele Rabosio

## Abstract

Analyses of pharmaceutical pipelines of drug development in the 1990-2010 documented progressively increasing attrition rates and duration of clinical trials, leading to a diffuse perception of a “productivity crisis”. We produced a new set of analyses for the last decade, using an extensive data of more than 45,000 projects between 1990 and 2017, and report a recent upsurge of R&D productivity within the industry. First, we investigated how R&D projects are allocated across therapeutic areas and found a polarization towards high-risk/high-reward indications, with a strong focus on oncology. Importantly, attrition rates have been decreasing at all clinical stages in recent years. In parallel, we observed an increase of early failures in preclinical research, and a significant reduction of time required to identify projects to be discontinued. Notably, more recent projects are increasingly based on novel mechanisms of action and target indications with small patient populations. Finally, by analyses of the relative contribution of different institutional types and development companies, we show that the observed increased performance in clinical trials is mostly due to the contribution of biotechnological companies, while pharmaceutical companies have significantly improved their performances in identifying false positives in preclinical research.

---

At the beginning of 2010s, many concerns were raised on the ongoing process of drug development, which culminated in a diffuse perception of a “productivity crisis” of the pharmaceutical R&D.^1, 2^ Data from the previous two decades showed a progressive increase of attrition rates throughout the whole pipeline of drug development and a significant increase of the time needed for the completion of clinical trials between the 1990s and the 2000s.^1, 3^

Several hypotheses were offered to explain these trends, including a gestation lag associated with the fundamental transformations of the scientific knowledge, underlying innovation dynamics within companies. Recently, however, fragmented signals have emerged of a change of tendency in the productivity of the drug development process: i) the number of New Therapeutic Entities (NTE) approved by year has been increasing since 2010;^4, 5^ ii) the oncology field has benefited from the extensive use of biomarkers;^8, 10^ iii) several innovations have entered pharmaceutical R&D, from artificial intelligence to aid decision-making throughout the process to 3D printing for personalized drug design and production.^9, 11^ In parallel, pharmaceutical companies are rethinking the entire R&D process, including the implementation of organizational solutions, such as the creation of “constellations” of dedicated centres^6^) and are devoting great efforts to devising strategies for more efficient early detection of the non-viable drug candidates.^7^ Finally, the recent upsurge of advanced therapies (e.g. CAR-T cell therapies for cancer treatment) has been interpreted as a sign of an ending gestation lag of further major breakthroughs.^8, 9^

Concurrently, regulatory agencies, such as the US Food and Drug Administration (FDA), have worked to accelerate drug approval. Requests for Breakthrough Therapy Designation,^12^ conceived to speed up the approval process for drugs that exhibit outstanding performances in preclinical research, have been increasing steadily since the onset of this initiative, passing from an average approval rate of 33% in its first years of application (2013-2015) to 44% in the last three years (2016-2018).

Here, using the same analytical approaches as in Pammolli et al., 2011,^1^ in order to ensure full comparability of results, we provide an updated and accurate picture of the current state of pharmaceutical R&D, using data on drug pipelines up to the 2018. Our data set includes more than 45,000 drug-development projects, whose processes have been registered with time and space signatures, potentially up to their marketing. Information on drug pipelines was integrated with links to an enriched patent database and to sales related to marketed compounds (between 2002 and 2016). These further refinements of the data allowed us to classify the indication associated with each project (i.e. to define the indication as “chronic”, “lethal”, “multifactorial” or “rare”) and to define the institutional type of project developer/originator.

We analyzed the therapeutic areas that are attracting a greater fraction of efforts by the pharmaceutical industry, and ascribed the observed changes in phase-by-phase attrition rates to specific institution types, using our classification of companies (i.e. pharmaceutical and biotechnological companies and non-industrial institutions). We used this classification to investigate the performance of different configurations of the division of innovative labor within the industry.^13^ Finally, we developed an indicator to monitor the evolution in time of the novelty of mechanisms of action associated with each project.

## Results

### The upsurge in pharmaceutical R&D productivity

We identified an R&D project as a specific indication-compound association, and selected projects started in either the US, Europe or Japan since 1990. We first focused on phase-by-phase attrition rates between 1990 and 2013 (Fig.1). At each stage, we defined a success when we observed a transition to the next stage of development within 4 years (that is why we considered only data up to 2013), or, in case of missing data, to any other subsequent phase, without time constraint. We used changepoint detection analysis^14^ to pinpoint the most relevant changes in linear regression slopes in the data and found that attrition rates in clinical phases have been declining in recent years, though they generally remained above their starting values. To portray a general picture of recent trends, we show in Table 1 the average values of phase-by-phase attrition rates in the three decades under study. As a general trend, we observed decreasing attrition rates in clinical trials, in recent years. Attrition rates in late-stages clinical trials (i.e. Phase II and III), however, remain quite high (Fig.1 and Table 1). Concurrently, failures in the preclinical phase have been increasing steadily, in conjunction with successful market launches (i.e. projects that are marketed after being registered by a regulatory agent, see the Registration panel in Fig.1).

**Figure 1:**
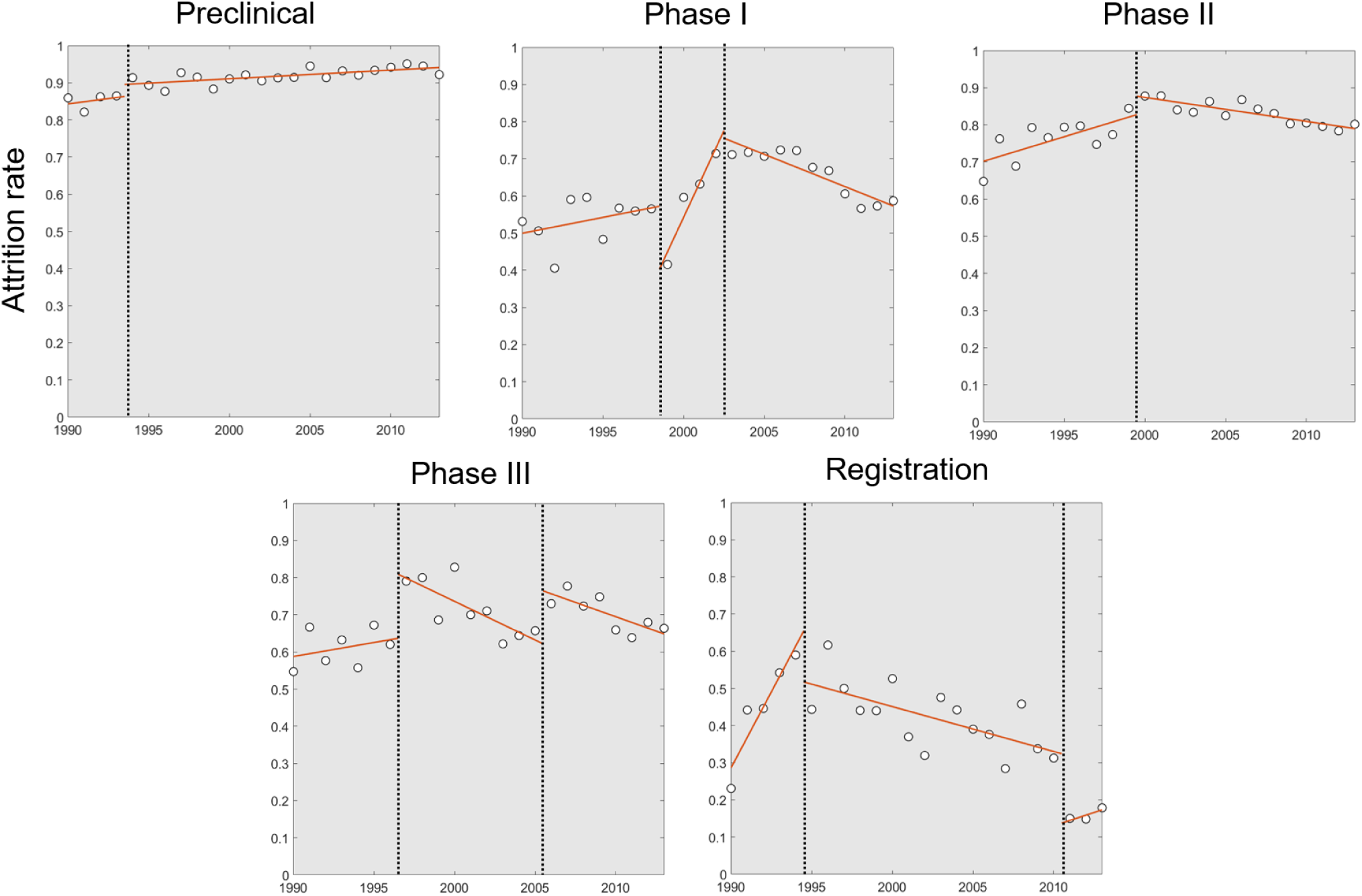
Attrition rates in time for different stages of drug development. White circles: data; red solid lines: linear regression in the corresponding time window; black vertical point line: changepoint. The attrition rate for a development phase in a year is defined as the percentage of projects that started the focal phase in that year and passed to the subsequent phase within four years (accordingly, 2013 is the last year we do consider).

**Table 1:**
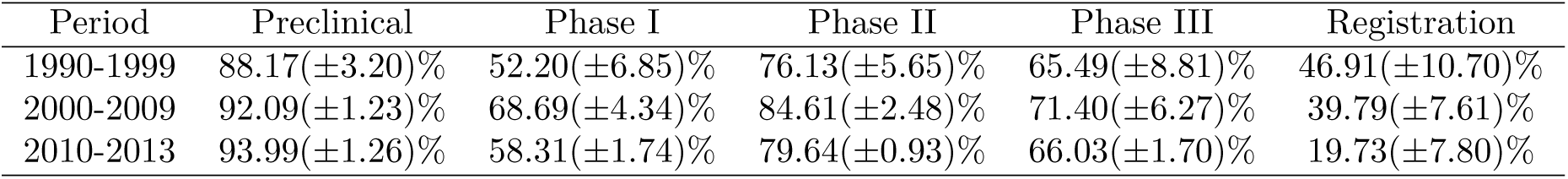
Average (± standard deviation) yearly phase-by-phase attrition rates in three different time intervals (1990-1999, 2000-2009, 2010-2013).

| Period | Preclinical | Phase I | Phase II | Phase III | Registration |
| --- | --- | --- | --- | --- | --- |
| 1990-1999 | 88.17($\pm 3.20$)% | 52.20($\pm 6.85$)% | 76.13($\pm 5.65$)% | 65.49($\pm 8.81$)% | 46.91($\pm 10.70$)% |
| 2000-2009 | 92.09($\pm 1.23$)% | 68.69($\pm 4.34$)% | 84.61($\pm 2.48$)% | 71.40($\pm 6.27$)% | 39.79($\pm 7.61$)% |
| 2010-2013 | 93.99($\pm 1.26$)% | 58.31($\pm 1.74$)% | 79.64($\pm 0.93$)% | 66.03($\pm 1.70$)% | 19.73($\pm 7.80$)% |

As a first attempt to identify drivers of decreasing attrition rates in clinical stages, we computed the relative performance of R&D projects targeting different therapeutic areas. To this end, we divided the projects according to their corresponding first-level Anatomical Therapeutic Codes (ATC), which consist of a hierarchical classification of drugs, maintained by the European Pharmaceutical Market Research Association (EphMRA). Fig.2 shows the relative contribution of each ATC class to the observed variations in phase-by-phase attrition rates, between 2000-2009 and 2010-2013. We computed the contribution of each ATC class as (Δ*AR*_*a,p*_ · *Sh*_*a,p*_)100*/*Δ*AR*_*p,tot*_, where: i) Δ*AR*_*a,p*_ is the observed *p*-specific attrition rate variation between the two periods for class *a*; ii) *Sh*_*a,p*_ is the share of phases *p* belonging to projects in class *a* in the period 2010-2013 and iii) Δ*AR*_*p,tot*_ is the total variation of the *p*-specific attrition rate between the two periods. Table S1 reports the values involved in this computation in their entirety (phase-by-phase attrition rates, project share, relative contribution by ATC class). Expectedly, the oncological ATC class L (Antineoplastic and Immunomodulating Agents; light blue in Fig.2) emerges as one of the most important in determining the observed variations, first and foremost because of its dominant share in almost all phases under consideration. While its contribution in absolute value is sizable in early development stages because of higher share, it becomes more significant in Phases III, where it is actually considerably higher than its share. In other words, the reduction of its attrition rate in Phases III has been more significant than in other ATC classes. In general, the contribution values did not deviate greatly from the project share distribution (Table 1), suggesting that most ATC classes have similar performances. In Phases III, instead, we observed a more diversified picture, with some ATC classes showing increasing attrition rates, while others significant decreases (the latter happens in particular for classes A: Alimentary Tract and Metabolism; J: General Anti-Infectives Systemic; L: Antineoplastic and Immunomodulating Agents and R: Respiratory System).

**Figure 2:**
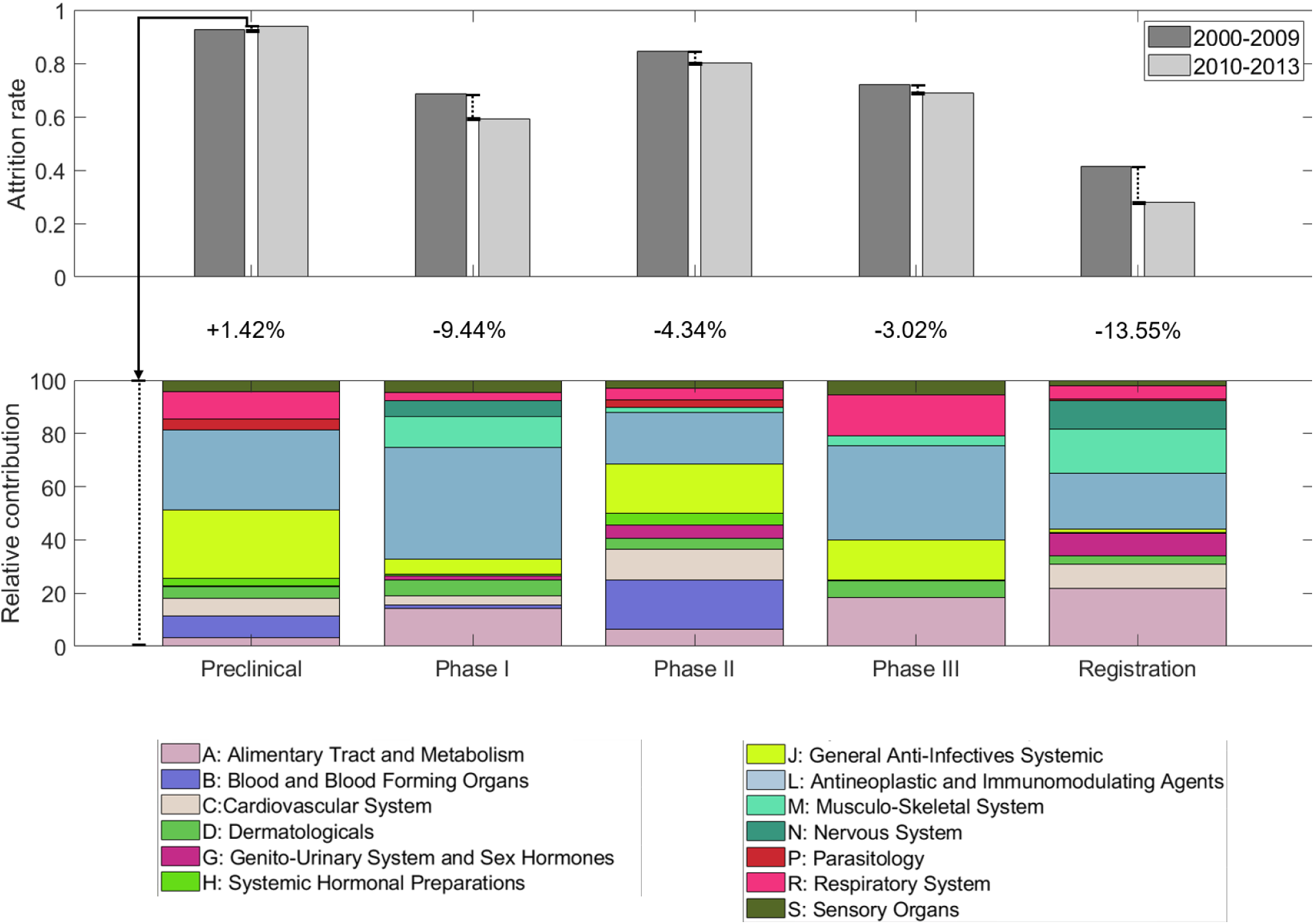
Variations in phase-by-phase attrition rates between 2000-2009 and 2010-2013 and the respective, relative contribution of each ATC class. Attrition rates in the above periods are computed as the total average attrition rate (i.e. 1 −*c*_*p,succ*,Δ*t*_*/c*_*p,tot*,Δ*t*_, where *c*_*p,succ*,Δ*t*_ is the total count of successful phases *p*, as defined in the text, in period Δ*t* and *c*_*p,tot*,Δ*t*_ is the total number of phases *i* in the same period. The relative contribution of its ATC class to the total variation in phase-specific attrition rates is computed as (Δ*AR*_*a,p*_ ·*Sh*_*a,p*_)100*/*Δ*AR*_*p,tot*_, where Δ*AR*_*a,p*_ is the observed *p*-specific attrition rate variation between the two periods for class *a, Sh*_*a,p*_ is the share of phases *p* belonging to projects in class *a* in the period 2010-2013 and Δ*AR*_*p,tot*_ is the total variation of the *p*-specific attrition rate between the two periods. Only positive contributions are taken into account in this computation. For complete values, see Table S1.

We also considered time as a fundamental variable. We first analyzed the relationship between the increasing attrition rate in the preclinical phases and the time needed to identify non-viable candidates. To this end, we looked at the time needed for discontinuation of projects and its distributions among dead projects in the two different time intervals (Fig.3). Interestingly, ≃ 70% of projects that had started between 2000 and 2009 were terminated in the year they entered preclinical research, with a ≃20% increase with respect to the previous decade. We then measured the time needed for a successful project to join the market. We choose patent application year as the starting time of each project. Accordingly, in the inset of Fig.3 we show the distribution of the time lag, measured in years, between patent filing and market launch of successful projects in the three different decades under study (based on the year of market launch). Interestingly, despite the increase observed between the 1990s and 2000s, this measure has decreased again, showing that the development of at least a fraction of the projects has become faster in recent years.

**Figure 3:**
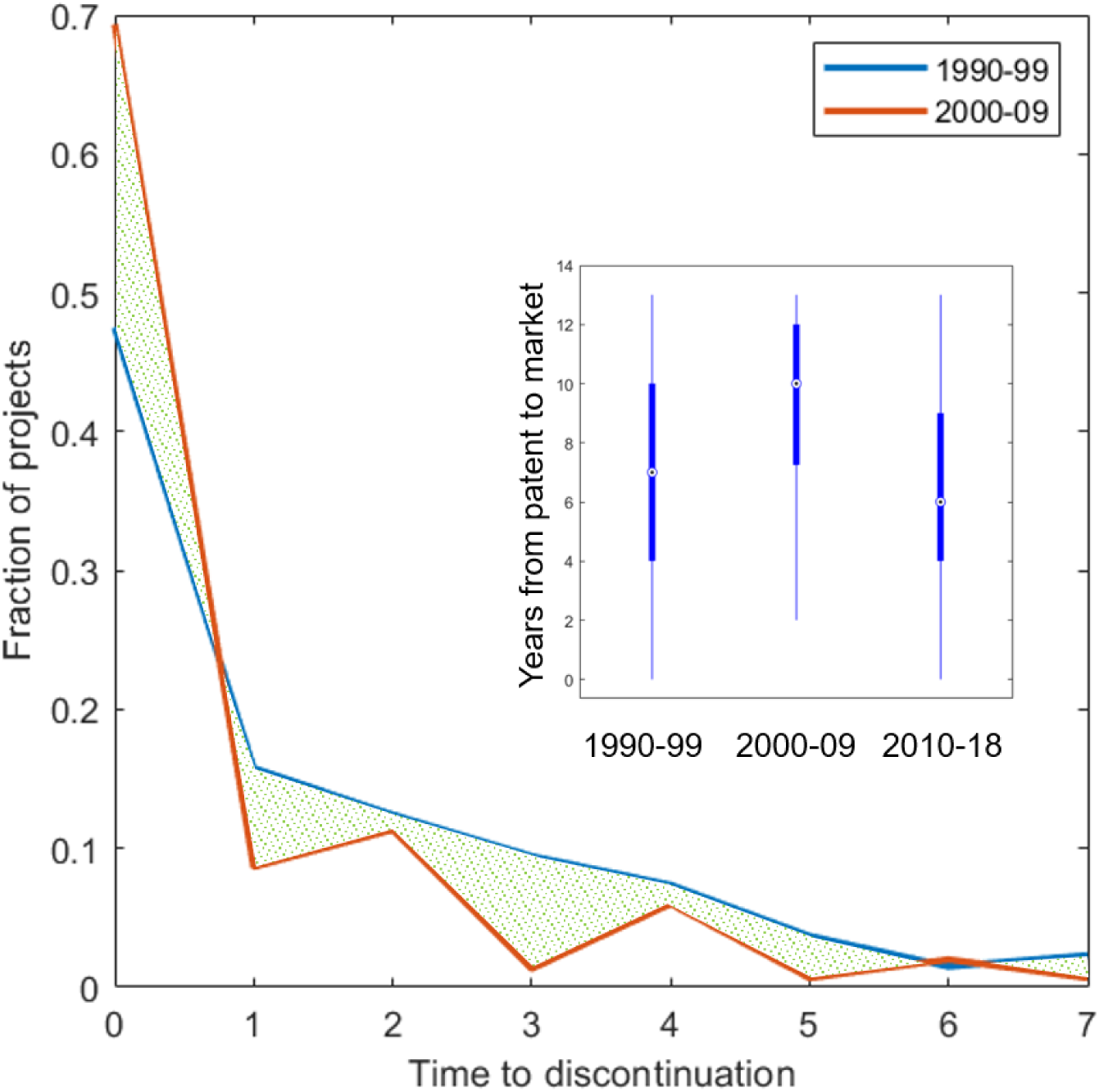
Time needed for project discontinuation in 1990-1999 (blue) and 2000-2009 (red). We highlight in green the area between the two curves. We show the fraction of projects that are discontinued after *x* years from the start of preclinical research. The distribution accounts for a maximum discontinuation time of 8 years, so we cannot perform such a computation for projects started after 2009. Inset: boxplot of the time interval between patent filing and market launch years, based on the year of market launch, in three different time intervals (1990-1999, 2000-2009, 2010-2013).

To track the evolution of phase duration, we computed the time needed to progress along the pipeline in the different decades of observation (in the 2010s, we limited the analysis to year 2013 and, to facilitate comparisons across decades, we imposed the constraint of 48 months as the maximum observable lag). As shown in Fig.4, the time needed to complete the preclinical phase is slightly increasing. When a project has entered clinical phases, its progress is instead signifcantly faster in Phase I, while the duration of Phase II has not increased, at least in the last decade. The duration of Phase III increased progressively in the analyzed decades and remains the longest, due to the complexity of inherent activities (the regulatory requirements, increasing patient sample size, the simultaneous multi-center logistics; see e.g. Scannell et al., 2012^2^).

**Figure 4:**
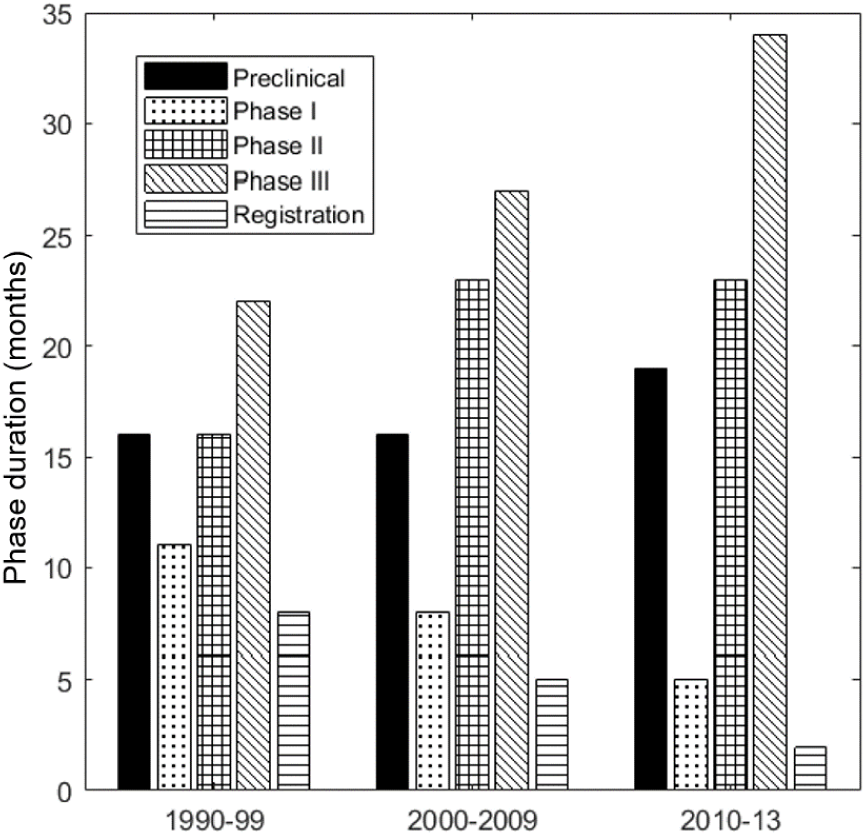
Median phase duration in time per each phase of drug development process, in three different time intervals (1990-1999, 2000-2009, 2010-2013). The duration of a development phase in a year is defined as the median time required to the projects that started the focal phase in the given year to pass to the subsequent phase. The median is computed considering only transitions with duration lower than or equal to four years, to make a sound comparison across decades.

### Finding the niche

All in all, the emerging picture is consistent with a general improvement of efficiency: attrition rates in clinical phases have all started decreasing in recent years, though as expected they remain higher than those at the start of the observation period. Moreover, there is significant evidence that the performance of early screenings has improved. Thus, we investigated which are therapeutic areas where the industry is focusing its efforts.

To this end, we partitioned the projects under study based on their ATC, identifying their main therapeutic areas at the 3-digit hierarchical level (ATC3). In Fig.5 we show how projects are distributed among therapeutic areas, as a function of the related probability of success (POS; i.e. how many projects reach the market from the preclinical phase, overall) up to 2013 and of the yearly average sales between 2002 and 2016. In general, results show that high-reward/high-risk projects (i.e. the area with low POS and high yearly sales) are polarizing the investments (Fig.5a), as previously reported.^1^ Expectedly, projects in therapeutic areas with higher revenues and lower attrition rates have registered the highest share increase between 2002-2009 and 2010-2017 (Fig.5b). In particular, monoclonal antibody neoplastics (class L1G) and immunosuppressants (L4X) have increased their share, while the still prevailing class L1X, being at lower POS and yearly sales, has remained constant.

**Figure 5:**
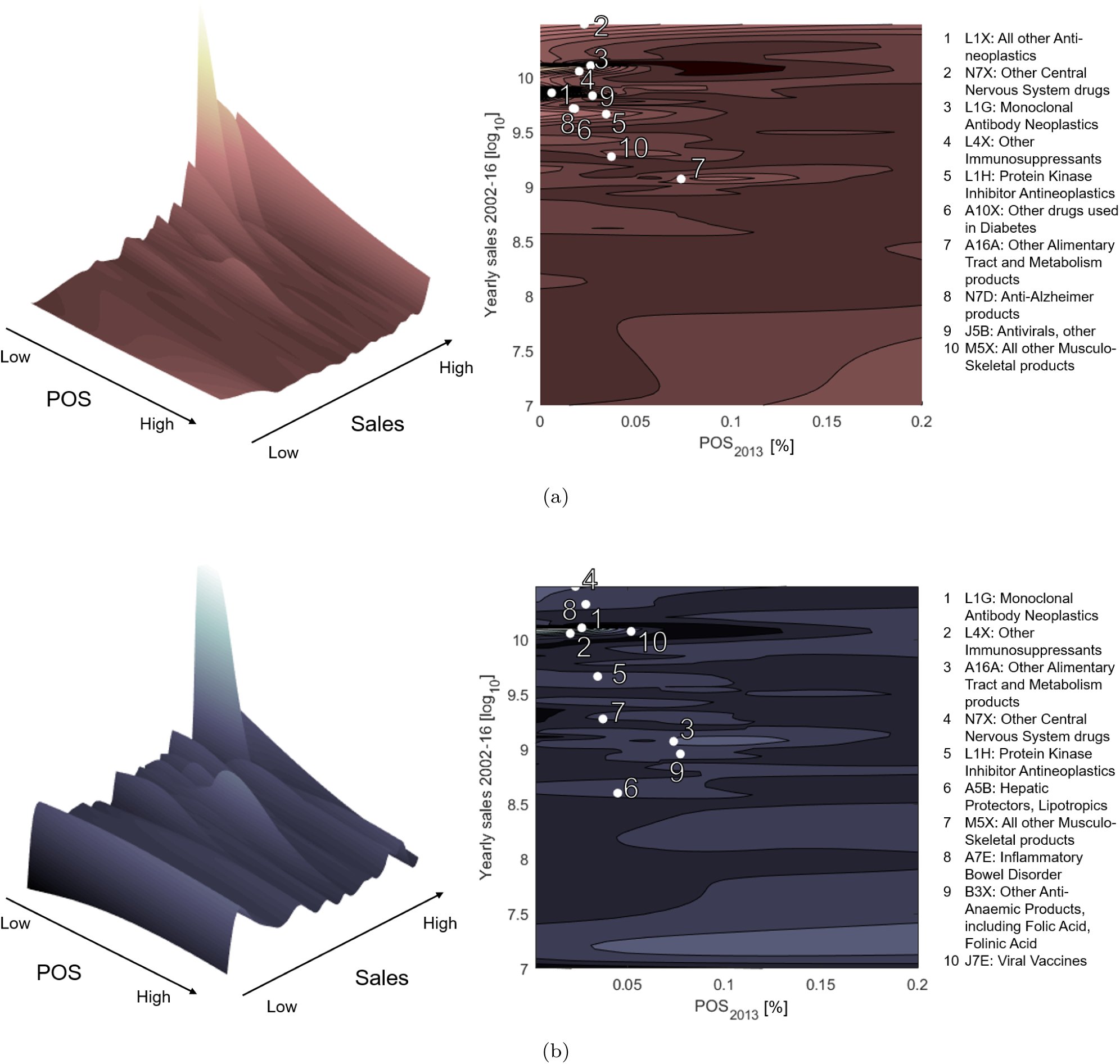
In each panel, the probability of success (POS) is shown on the x axis and the logarithm of potential sales (yearly average computed in 2002-2016) on the y axis. A contour plot and a three-dimensional view of the same distribution are shown. In the contour plot we highlight the top 10 ATC3 classes by the focal metric being shown on the vertical axis. These are listed besides the contour plots. a: The vertical axis shows the percentage distribution of research and development (RD) projects by POS and potential sales level. The distribution of RD efforts is concentrated in the upper left hand corner of the plot (indicating high sales and low POS); b: The vertical axis shows the share variation between 2002-2009 and 2010-2017, again as a function of POS and sales.

The concentration of projects in oncology is even more apparent in the latest decade: i) 4 out the top 5 ATC3 classes, ranked by their overall share in 2010-2017 projects, are related to oncology (L category); and ii) more than 40% of ongoing studies, currently listed on ClinicalTrials.gov^1^, are oncology-related (see Table S2). Recent reports^5, 11^ showed similar findings and predicted even greater sales and market share for the future for oncological treatments. The observed concentration of projects in oncology can be related to the increase of the R&D productivity, as oncology is recognized among the areas with higher unmet medical need and where major improvements can still be made.^2^ In this respect, we found that advanced therapies (i.e. cell or gene therapies) are mostly dedicated to oncology (Fig.S1), and new projects have been on a steep rise in the last few years (Fig.S2), as recently reported.^9^ Also, the rising importance of anti-cancer antibodies (class L1G) is a factor of simplification of drug preparation for preclinical test and clinical trials, as the efficiency of monoclonal antibody production has significantly improved in recent times.^21^ Other relevant fields that showed up in rankings include degenerative diseases of the central nervous system (N7X), with specific reference to Alzheimer’s disease (N7D), another area in which medical need is high.^16, 17^ Interestingly, while projects in class L have improved their attrition rates after 2010 (see Fig.2 and Table S1), possibly due to the drivers we just mentioned, the performance of projects in class N worsened in most cases (i.e. fewer failures in preclinical research, and more in late clinical development stages, Phase II and III). In fact, IQVIA (2019)^9^ reports that, out of 86 projects on Alzheimer’s disease in the last ten years, only one received approval.

The progressive focus of the industry on projects of high complexity, in relatively unexplored areas, can also be seen horizontally, across therapeutic areas. Orphan drugs approval, for instance, has significantly increased over recent years.^18^ The number of yearly NME approvals for orphan drugs has more than doubled from 2000-2009 to 2010-2017, while drug repositioning approvals towards rare diseases have tripled in the same period.^18^ We used a manual classification of indications to retrieve the share of rare diseases (defined as having a prevalence of ≤200,000 affected individuals in the US) by year of project start (Fig.6a). In the observation period, this share has increased from 3% in 1990 to about 16 % in 2017, suggesting that the greater focus on orphan drugs might be an important driver also in the improved efficiency of the whole R&D process. On one side, FDA data^19^ show that orphan drugs cover a majority share in fast-track programs. On the other, the “better than the Beatles” problem described by Scannell et al. (2012)^2^ might be less relevant for these diseases. We report in Table S3 that the average phase-by-phase attrition rates have also been declining in the subset of projects focusing on rare indications, with the notable exception of Phase III, in which trial set-up is known to be more demanding because of the limited size of target population.^20^

**Figure 6:**
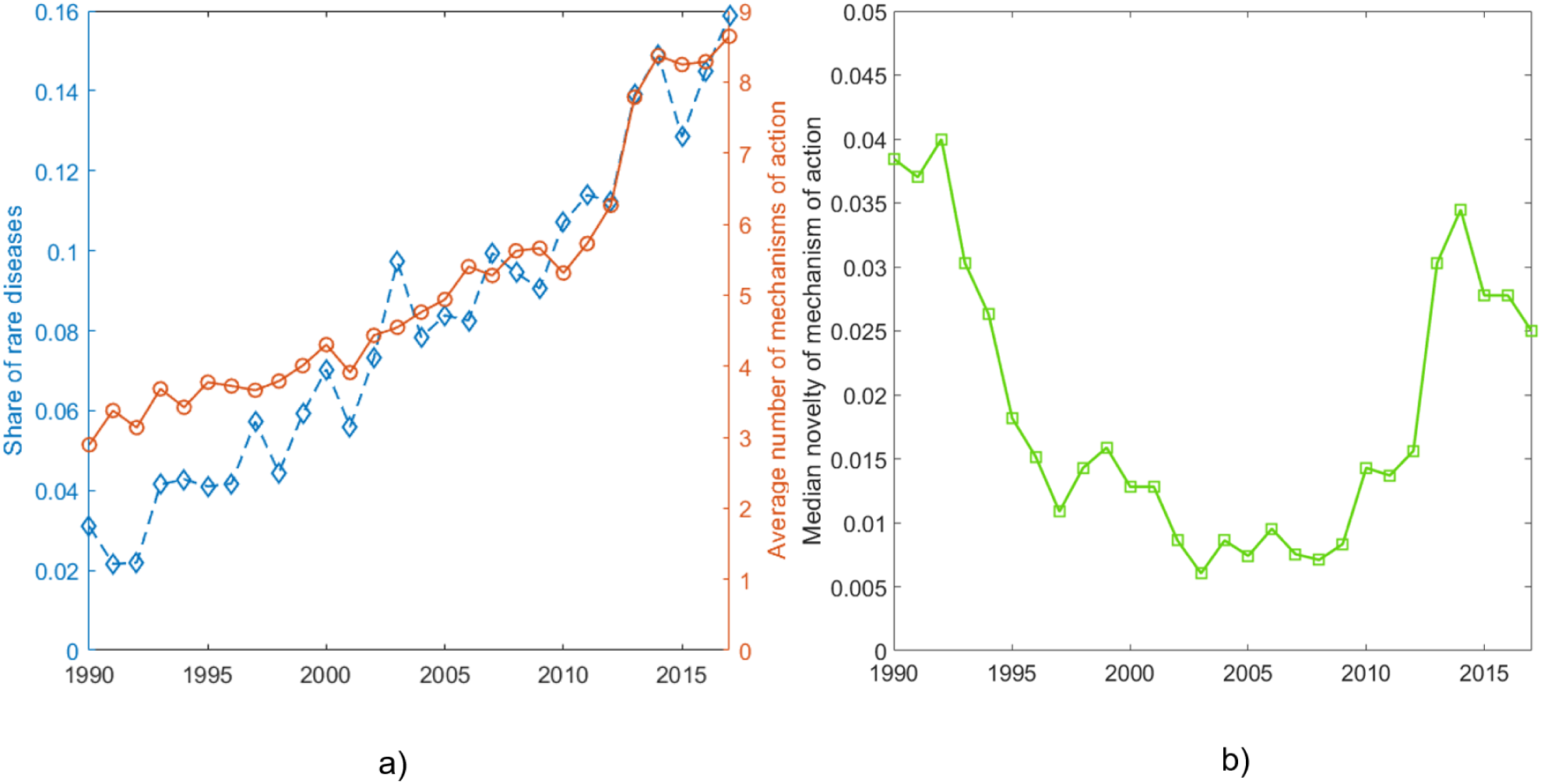
a: Evolution in time of the share of projects targeting rare diseases (i.e. having a prevalence of fewer than 200,000 affected individuals in the US) and of the average number of mechanisms of action per project, between 1990 and 2017, by project starting year (i.e. the year the focal project entered preclinical research). b: Evolution in time of median novelty of mechanism of action per project, between 1990 and 2017, by project starting year.

Expectedly, the general tendency towards orphan drug development is also a factor of increasing difficulty of projects. It has been observed, for instance, that orphan drug development takes, on average, 2.3 years longer.^20^ This is due, for instance, to the smaller and geographically dispersed patient populations and to the scarcity of available animal models and biomarkers. An additional factor of complexity might be the increasing relevance of multifunctional drugs, which have emerged, in opposition to single-targeted drugs, as a new approach to treatment^22, 23^ (see Barabasi et al., 2011^25^ on the concept of network medicine). We also considered the average number of mechanisms of action per drug, by the starting year of the project (Fig.6a). Also, here we observed a clear positive trend, with an even more pronounced increase after 2010. Between 1990 and 2017, this number has nearly tripled. This may reflect a general improvement of drug efficacy, as they act on multiple targets, or, alternatively, it may represent the increasing difficulty of drug design.

Finally, we assessed the degree of novelty of this increasing number of mechanisms of action. To this end, we devised an indicator that measures the number of times a given mechanism of action, listed in the focal project, has appeared in projects started previously, taking into account the total number of previous projects (see Methods for details). Interestingly, the median value of novelty of mechanism of action in projects, despite decreasing from 1990 to 2010, has then started to increase again, overcoming the obvious bias imposed by the increasing probability of finding previous projects with the same mechanism of action (Fig.6b). Recent reports^9^ showed, in fact, that 34% of mechanisms of action in FDA-approved drugs in 2018 were first-in-class (i.e. they were different from those of existing therapies). To gain insights into the effect of novelty on project success, we divided our dataset in successful (i.e. marketed) and failed projects and found a significantly higher median novelty of successful projects (0.083 vs 0.015; a Wilcoxon^24^ test rejects the null hypothesis that the two distributions have the same median with p«0.01).

### The Division of Innovative Labor

Having assessed a trend inversion for phase-by-phase attrition rates, we then endeavoured to ascribe these patterns to specific institutional categories. As shown in Pammolli et al., (2011)^1^ and Arora et al. (2009),^26^ the contribution of different institutional categories (pharmaceutical and biotech companies or non-industrial institutions) to R&D performance might differ significantly. To investigate their role in the observed increasing productivity, we used a manual classification of a large number of companies, according to the main institutional types (i.e. pharmaceutical, biotech, university, hospital, other research center). We extended this classification to the list of developer institutions in our data set. Overall, this classification allowed us to classify ≃60% of projects, that showed statistics comparable to the whole sample (Table 1). In Fig.7 we show the percent contribution of each type of institution to the change we observed in each phase, while in Table 2 we list the complete results concerning the phase-by-phase attrition rates for each institution type and the corresponding project share, in the two periods. We measured the contribution of institution type *i* to the variation in attrition rates in phase *p* between 2000-2009 and 2010-2013 using the formula *δ*_*ip*_ = (Δ*AR*_*ip*_ *Sh*_*i*_) *100*/*Δ*AR*_*tot,p*_, where Δ*AR*_*ip*_ is the variation observed in attrition rates in *p* in the projects developed by *i, Sh*_*i*_ is the share of phases belonging to projects developed by *i* in 2010-2013, and Δ*AR*_*tot,p*_ is the total attrition rate observed for phase *p* for all projects for which institution classification was available. Results showed that project phase-by-phase share alone is not sufficient to account for the percent contribution to the attrition rate change, suggesting that some minority institution types have greatly improved their performance. For example, despite pharmaceutical companies have a lower prevalence of projects in preclinical research (Table 2), their contribution to the increased attrition rate in that phase is predominant (Fig. 7). Conversely, the contribution of biotech companies is greater than its share in late clinical stages (Phase II and III).

**Table 2:** Average phase-by-phase attrition rates and phase-by-phase share in 2000-2009 (00) and 2010-2013 (10), for the three institutional types under study (pharmaceutical and biotechnological companies and non-industrial institutions).

| Developer | Attrition rates |  |  |  |  |  |  |  |  |  |
| --- | --- | --- | --- | --- | --- | --- | --- | --- | --- | --- |
| | Pr | | $P_I$ | | $P_{II}$ | | $P_{III}$ | | Reg | |
|  | 00 | 10 | 00 | 10 | 00 | 10 | 00 | 10 | 00 | 10 |
| Pharmaceutical | 87.95 | 92.59 | 70.16 | 62.18 | 79.34 | 76.90 | 65.07 | 62.53 | 40.95 | 20.00 |
| Biotech | 89.68 | 90.16 | 63.95 | 55.77 | 84.50 | 78.10 | 77.40 | 67.42 | 31.68 | 19.75 |
| Non-industrial | 96.61 | 96.43 | 66.67 | 56.56 | 87.02 | 75.00 | 81.36 | 58.82 | 48.65 | 0 |
| Developer | Share |  |  |  |  |  |  |  |  |  |
| | Pr | | $P_I$ | | $P_{II}$ | | $P_{III}$ | | Reg | |
|  | 00 | 10 | 00 | 10 | 00 | 10 | 00 | 10 | 00 | 10 |
| Pharmaceutical | 26.35 | 26.98 | 43.54 | 48.32 | 46.74 | 47.84 | 62.46 | 62.17 | 77.08 | 59.09 |
| Biotech | 48.49 | 47.38 | 45.46 | 43.33 | 43.45 | 44.61 | 31.23 | 32.79 | 16.78 | 36.82 |
| Non-industrial | 25.16 | 25.65 | 11.00 | 8.35 | 9.81 | 7.55 | 6.31 | 5.04 | 6.51 | 4.09 |

**Figure 7:**
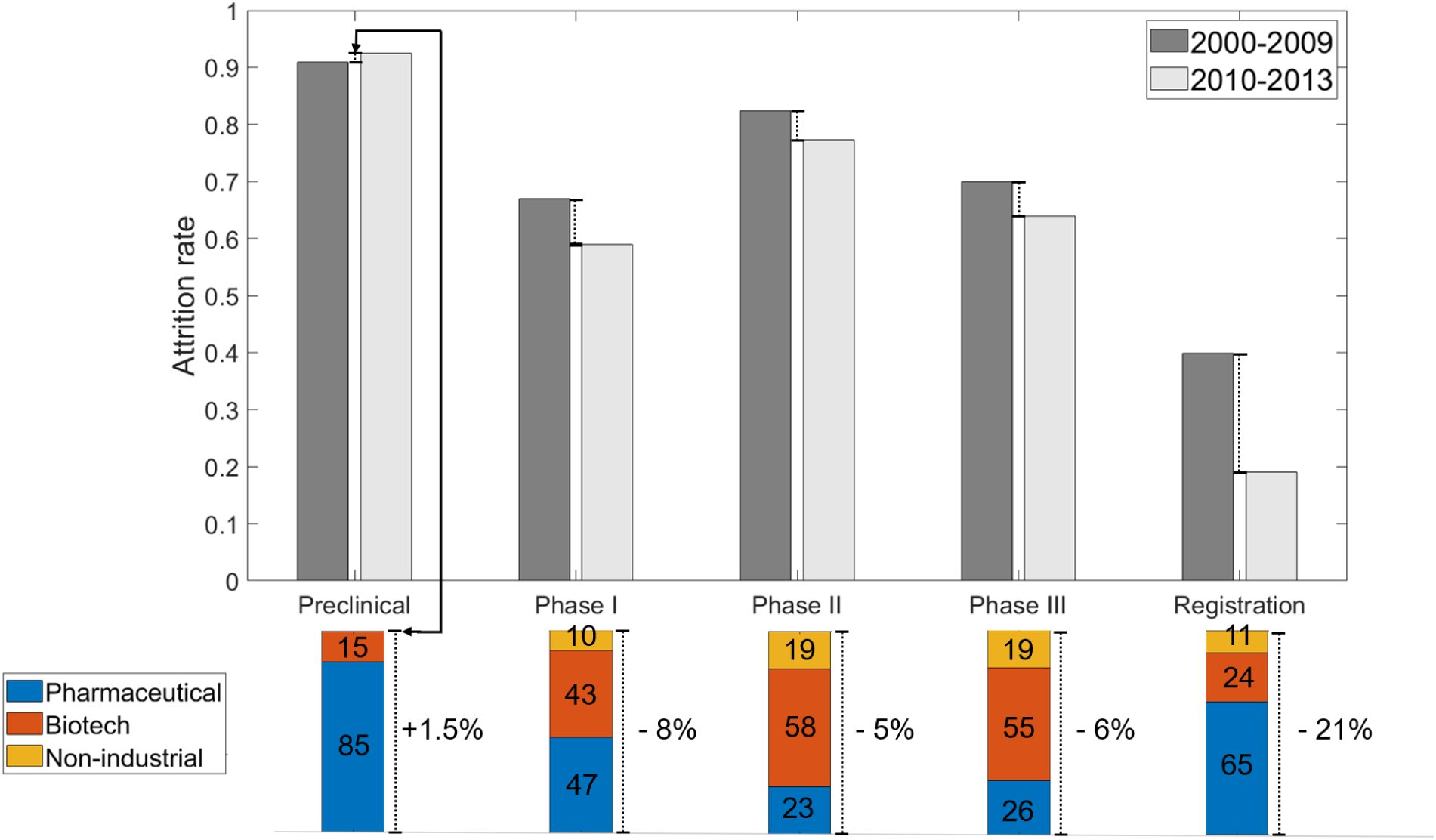
Contribution to each of the three main institutional categories to changes in global attrition rates between the periods 2000-2009 and 2010-2013. In the upper bar plot we show phase-by-phase attrition rates in the two periods. The bars below expand the attrition rate variation for the focal phase and are divided according to the relative contribution of each institutional type to the observed variation.

Moreover, by using the same classification of institutions, our database allowed us to define different Originator-Developer relationships, in which originators are defined according to the patent assignees we have retrieved from our linked patent dataset. Again, division of labor might unravel differences in R&D performance and help to define a landscape of drug development organization in greater detail, as, in general, R&D agreements in the pharmaceutical industry have become increasingly important.^27^ For instance, academic and non-industrial institutions have been advocated as pivotal in driving early development of candidate drugs.^28^ Other sources^29, 30^ underline the importance of interfirm and public/private knowledge transfer to influence R&D productivity. We studied the effect of different Originator-Developer (OD) relationships on the productivity indicators we have analyzed so far, namely attrition rates and sales (i.e. the logarithm of composite sales in the available period for sales – 2002-2016). In general, we identified an OD relationship for 4968 projects in the selected time span (1916 in the decade 1990-1999, 3052 in 2000-2013). We show in Table S5 a full count of these projects by their relative OD relationship. We found that biotechnological companies have increased their share after 2000 as either originators or developers, while pharmaceutical companies are no longer dominant. This trend is confirmed by recent reports.^9^ In particular, the biotech–biotech relationship has increased its share.

Taking the pharmaceutical – pharmaceutical relationship as our baseline OD relationship, we show in Table 3 the results of the regressions accounting for different OD relationships against the baseline (the complete results can be found in Tables S6–S10). We repeated the regression for data before and after 2000, taken as a reference year (projects starting before 1990 are not accounted for). We also considered fixed effects of time and project difficulty (expressed by indication and by our indication classification, “chronic”, “lethal”, “rare” or “multi-factorial”).

**Table 3:**
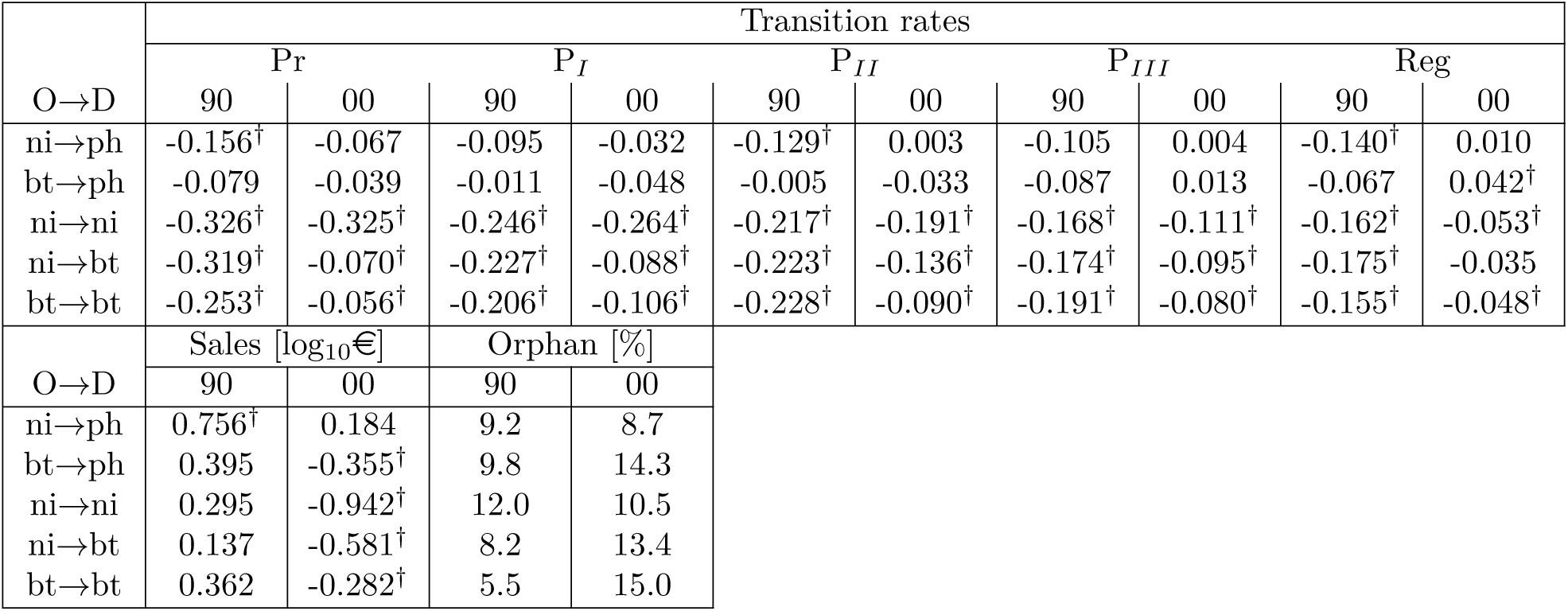
Regression coefficients for the five cases of Originator-Developer relationship and the three categories of response variables (phase-by-phase transition rates, phase duration and sales). *ni*: non-industrial; *ph*: pharmaceutical; *bt*: biotech. †: significant at *p* < 0.05.

A few observations, arising from the results shown in Table 3, are worth mentioning. First with respect to the role of non-industrial originators in the drug development process, we did not find a significant difference in the share of projects with a non-industrial partnership between projects started in the 1990s and after 2000 (both being around 20%). We found, instead, that the non-industrial → pharmaceutical model has changed its performance considerably before and after 2000. In the former case, we found evidence that attrition rates were higher than the pharmaceutical → pharmaceutical baseline. In the end, though, projects turned out to be significantly more successful (i.e. to have greater sales with respect to the baseline) on the market. Notably, in the latest period, we found that differences from the baseline model are no longer tangible. In parallel, an obvious change is affecting the non-industrial → non-industrial model. We found evidence that attrition rates remain high (or higher, even) in early development phases, but they are also decreasing in later stages, with respect to the baseline.

A few interesting remarks come from the analysis of the evolution of the performance of biotechnology firms. In the biotech → pharmaceutical model, we did not find significant differences from the baseline and in the two time periods, except for the fact that the effect on sales has changed sign after 2000. This is a pattern that can be found in all other OD relationships including biotech firms as either originators or developers. For the non-industrial biotech → model, the considerations for the strictly non-industrial model remain valid, but with a more drastic reduction of the difference in attrition rates with respect to the baseline after 2000 (and no sensible change in terms of phase duration, though).

The general pattern of sales becoming significantly lower in projects involving biotech firms or non-industrial institutions, after 2000, hints at a possible greater focus on smaller markets of this part of the industry. For this reason, we list in Table 3 the share of projects focusing on an indication classified as “rare” in our expert-guided classification. Preliminarily, we report that the pharmaceutical baseline model increased its share from 4.5% to 10.7%. In the first place, we observed that projects involving non-industrial institutions, in any role, have not increased their share, quite the opposite. Also, they appear, after 2000, comparable to what is observed in the baseline. At the same time, we found that projects in which non-industrial institutions appear among originators and developers both show the greatest difference in sales with respect to the baseline, almost one order of magnitude. This result reveals, possibly, an increasing focus on rare indications and on small target populations. Concurrently, we found that the biotech → biotech model has had the greatest increase in orphan share in the two periods, including the baseline.

## Discussion

The sustainability and productivity of R&D on drugs remains a major issue for the biopharmaceutical industry. Our analyses of drug development pipeline data, though, showed significant improvements in the overall efficiency of the process. In particular, the screening of drug candidates is becoming more and more effective in the selection of more viable projects at early stages, so that later attrition rates are decreasing. This has a two-fold effect on productivity: increased approval of drugs and decreased average cost per project. As a possible driver of this phenomenon is the increased attention to the validation of the drug targets in preclinical research, both in terms of their role on the disease, and the toxicity inherent to their manipulation. Indeed, the extensive genetic validation of drug targets has been shown to improve the chances of passing through clinical stages^32^ and has become more widely embraced in different therapeutic areas.^33, 34^ A better selection of patient subsets for the clinical trials via “stratification” based on biomarkers^36^ is another possible factor of improvement of success rates. Also, the prevalence of antibodies as new drugs has simplified both preclinical development and clinical grade batch preparation, since antibody production is a relatively easy process amongst biologics. Finally, “precision” diagnostic assays have been increasingly used as clinical endpoints,^35^ thus providing accuracy and efficiency to clinical trial assessment.

We found that many of these improvements are quite widespread across projects in different therapeutic areas and at different stages of clinical development, except for Phase III, in which performances can still vary greatly and where molecular stratification of patients is still very poor. Also, we found that the pharmaceutical industry is increasingly focusing on therapeutic/pathological areas where medical need is high (i.e. oncology and degenerative diseases of the CNS). These increasing efforts on high-risk projects, combined with the rise in the number and novelty of mechanisms of action in recent projects, reveal that new discovery areas are opening up, in a renewed pursuit of the “endless frontier”. We found, though, that phase duration in late stages of drug development is consistently increasing, particularly in Phase III, pointing at an increasing difficulty of project requirements in terms of trial characteristics and efficacy. Though a recent publication reports similar findings,^9^ it speculates that an increasing focus on orphan drugs may also result in shorter trial duration. We documented instead further evidence of increasing numbers of drugs based on disease mechanisms, which follows improvements in the understanding of the etiology of diseases. Though this paradigm shift may result into the generation of more efficacious drugs, it might affect the length of the process of drug design, as shown by the duration of successful preclinical research that, in fact, has increased after 2010.

We also found that the intensification of the collaboration between firms and regulatory agents can guide the whole process from the beginning and positively impact the development time in Breakthrough Therapy Designation procedures, as recently suggested.^31^

When looking at the relative contribution of different institutional types to the increasing R&D productivity, we found that results depend on the focal phase of drug development. The observed increase of early failures, i.e. in preclinical research, are mostly attributed to pharmaceutical companies. In late stages of clinical trials (Phase II and III), we found that the performance of projects developed by biotech companies has documented the most significant increase. Non-industrial projects are also contributing more than their (minority) share to the improvement of attrition rates in late clinical phases. Regarding the institutional models regulating the originator-developer scheme, projects originated and developed by non-industrial institutions (i.e. universities, hospitals and other research centers) are found to fail earlier, when started after 2000, than in the previous decade, and they reduce the duration of Phase II and III, the most critical for R%D costs, in a statistically significant way. Concurrently, we found that collaborations between non-industrial institutions and pharmaceutical companies, which were enablers for state-of-the-art technology development in pharmaceutical firms (something we found in our data in the 1990 decade and is reported in the literature, see e.g. Cassiman & Gambardella, 2009^37^), no longer retain the same signature after 2000. Additionally, this is not due to an increasing focus on the orphan drug market, which might overshadow the added value of knowledge outsourcing. On the other side, we found evidence that non-industrial institutions are focusing on extremely rare diseases, as sales related to strictly non-industrial projects are lower than the baseline by nearly an order of magnitude, after 2000. We did find, also, that projects involving biotechnological firms have all increased their share of projects on rare diseases, and they appear higher than the baseline by 3-5% after 2000.

## Methods

### Data

The main data source employed in this study is R&D Focus, a proprietary database about drug R&D projects. Moreover, R&D Focus data has been complemented with proprietary sales information from the IQVIA sales database, and with free patent data from Regpat and USPTO datasets.

R&D Focus contains information about over 43,000 medical compounds developed until September 2018, both successful and failed. For each compound a number of details are available. In particular, in this work we use the following pieces of information:

- ATC codes, dividing compounds into groups on the basis of the organ on which they act and their therapeutic and chemical properties; it is a hierarchical classification envisaging five levels, from ATC1 to ATC5.
- Indications, i.e., the diseases for which the compound is/will be used. We can also take advantage of a classification of the indications provided by a pharmacologist; the indications are classified as rare/not rare, lethal/not lethal, chronic/not chronic, multifactorial/not multifactorial. A disease is multifactorial when its causes are represented by the competition of several factors of a different nature, apparently not in direct connection between each other.
- Mechanisms of action, representing the biochemical interactions through which the drug produces its effects.
- Companies which have developed the compound.
- Codes of the patents possibly related to the compound.

Moreover, each pair (compound, indication), which for us defines a project, is connected with information about its development history. The development history is the sequence of development phases that the compound has undergone until its marketing or failure for that indication; the available phases are Preclinical, Phase I, Phase II, Phase III, Registration, Marketed. Each phase in the history is associated with date and country; in this work only the projects initiated in USA, EU or Japan are taken into account. Overall, this selection and processing and data originates 49,591 projects.

The IQVIA sales data at our disposal envisage the sales in euros of both branded and unbranded pharmaceutical products from 2002 to 2016 in 35 countries. The database contains 202,651 products corresponding to 48,402 distinct compounds. The compound names in IQVIA have been linked to the R&D Focus compounds via text matching. In R&D Focus there are 2,333 marketed compounds, and we have been able to connect 2,123 of them with IQVIA sales entries (91.0%).

The patent codes specified in R&D Focus are used to link the compounds with the freely available patent datasets (the American USPTO dataset, and Regpat for EPO and PCT patents); the R&D Focus compounds are associated with 2,917 USPTO patents, 3,441 EPO patents and 2,419 PCT patents. The integration with the free datasets allowed us to retrieve the assignees of the patents.

The assignees of the patents, as well as the companies developing the compounds specified in R&D Focus, are divided into some categories through a manual classification provided by a domain expert; in particular, we consider three industrial categories (pharmaceutical, biotech and other industrial) and three non-industrial categories (university, hospital and other research centers). About the coverage of the manual classification, 84.6% of the R&D Focus patents have all the assignees classified, and 87.7% of the R&D Focus compounds have all the developing companies classified.

## Data processing

### Attrition Rate

The attrition rate for a development phase in a year is defined as the percentage of projects that started the focal phase in the given year and passed to the subsequent phase within four years (accordingly, the maximum possible starting year in our data is 2013). If the following phase is missing but a more advanced one is recorded, then the transition is deemed accomplished without imposing time constraints.

### Phase duration

The duration of a development phase in a year is defined as the median time required to the projects that started the focal phase in the given year to pass to the subsequent phase. The median is computed considering only transitions with duration lower than or equal to four years, to make a sound comparison across decades.

### Probability of success

We measure the probability of success of projects associated with a specific ATC3 as the number of projects which reach the market (i.e. they have a “Marketed” phase in their history) over the total number of projects in that ATC3. In this paper, we do not take into account projects started after 2013.

### Novelty of mechanism of action

We express the degree of novelty of mechanisms of action of a project as:

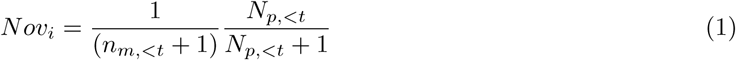

where *n*_*m*,<*t*_ is the minimum number of times a mechanism of action listed in project *i* has appeared in previous projects, while *N*_*p*,<*t*_ is the total number of previous projects.

## Statistical techniques

### Changepoint Detection

Changepoint detection^14^ identifies the time instants (changepoints) corresponding to abrupt changes in a function. Identifying the changepoints divides the function into sections, and in particular we split the attrition rate in correspondence of the years where the regression line changes the most. This is obtained by finding the sections of the function such that the sum of the residual errors of the regressions in each section is minimized.

Note that adding more changepoints keeps reducing the value of the residual error, leading to over-fitting. To avoid this problem, the error metric needs to envisage also a term penalizing high number of changepoints.

Let *x*_1_, *…, x*_*n*_ be the points of the function that we are studying, and let *f* ^*p,q*^ be the regression line approximating the function between the time instants *p* and *q* (*p* < *q*). The changepoint detection procedure finds the *K* time instants *k*_1_, *…, k*_*K*_ minimizing the following metric:

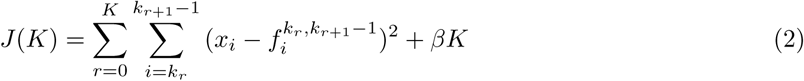

where in this formula *k*_0_ represents time instant 1 and *k*_*K*+1_ represents the last time instant (*n*). The internal summation describes the residual error of the regression between the time instants *k*_*r*_ and (*k*_*r*+1_ − 1). The term *βK*, where *β* is a parameter to be set, penalizes the addition of new changepoints. It can be easily shown that a new changepoint is rejected if it does not bring an improvement to the residual error of the regression at least equal to *β*. In this work the threshold *β* has been set to twice the variance of the function, meaning that we stop adding changepoints when the subsequent new one would increase the *R*^2^ determination coefficient of the regression of less than 2*/n*.

### Regression with dummy variables

We model here a set of response variables in a regression framework:

- phase-by-phase transition: binary variable identifying the successful passage from the focal phase to the next one;
- sales: logarithm of the sum of sales of the focal drug.

Our main explanatory variable is the binary variable identifying the type of Originator-Developer (OD) relationship under study. We define the originators of a project according to the assignees of the related patent(s), and the developers according to the developing company listed in the relevant R&D Focus project. We define OD relationships according to the presence of at least 1 assignee or developer in one of the relevant institutional types. We treat the “university”, “hospital” and “other research” classifications as “non-industrial”. Also, we define as the baseline OD relationship the one that has a pharmaceutical company among originators and developers both. Then, we study 5 possible relationships: non-industrial (O) and pharmaceutical (D); biotech (O) and pharmaceutical (D); non-industrial (O) and non-industrial (D); non-industrial (O) and biotech (D); biotech (O) and biotech (O). We treat each project which has at least one non-industrial institution among originators and developers as a non-industrial-non industrial project (the same goes for biotech firms). Mixed projects (but missing the focal type of institution in either the originators or the developers) are included in the baseline.

In addition, we use a few dummy variables to control from fixed effects characterizing the focal project: the starting year, the indication and the classification of the indication along four dimensions (i.e. “chronic”, “lethal”, “rare” and “multi-factorial”). The latter has been performed manually by a domain expert.

It follows that the regression model for the generic response variable *X* can be written as:

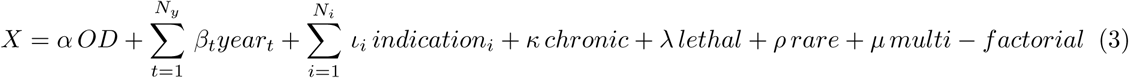

where OD is the binary variable classifying each project by either a relevant project according to the OD relationship under study, or a baseline project (pharmaceutical as originator and developer both).

---

**Table S1:** Average phase-by-phase attrition rates in 2000-2009 (00) and 2010-2013 (10), phase-by-phase share in 2010-2013, and relative (positive) contribution, when projects are divided according to their first-level ATC class. The relative contribution of its ATC class to the total variation in phase-specific attrition rates is computed as (Δ*AR*_*a,p*_ · *Sh*_*a,p*_)100*/*Δ*AR*_*p,tot*_, where Δ*AR*_*a,p*_ is the observed *p*-specific attrition rate variation between the two periods for class *a, Sh*_*a,p*_ is the share of phases *p* belonging to projects in class *a* in the period 2010-2013 and Δ*AR*_*p,tot*_ is the total variation of the *p*-specific attrition rate between the two periods. Only positive contributions are taken into account in this computation.

| ATC class | Attrition rates |  |  |  |  |  |  |  |  |  |
| --- | --- | --- | --- | --- | --- | --- | --- | --- | --- | --- |
| | Pr | | $P_I$ | | $P_{II}$ | | $P_{III}$ | | Reg | |
|  | 00 | 10 | 00 | 10 | 00 | 10 | 00 | 10 | 00 | 10 |
| A | 88.49 | 88.99 | 68.54 | 55.92 | 79.11 | 76.86 | 67.95 | 61.72 | 49.28 | 26.67 |
| B | 89.37 | 93.62 | 57.69 | 52.94 | 81.92 | 57.45 | 74.07 | 76.27 | 35.90 | 63.64 |
| C | 92.71 | 94.58 | 69.85 | 62.30 | 86.09 | 77.96 | 74.68 | 74.74 | 44.21 | 18.52 |
| D | 87.77 | 91.84 | 63.64 | 41.82 | 89.11 | 78.61 | 61.80 | 68.67 | 40.63 | 34.21 |
| G | 84.72 | 85.00 | 71.10 | 66.27 | 85.98 | 80.92 | 68.29 | 67.86 | 48.24 | 31.58 |
| H | 87.38 | 100 | 79.59 | 70.59 | 79.17 | 61.29 | 60.00 | 64.29 | 29.41 | 25.00 |
| J | 91.45 | 93.95 | 65.46 | 59.62 | 83.70 | 73.91 | 68.61 | 61.11 | 30.08 | 28.30 |
| L | 95.55 | 96.58 | 67.02 | 58.73 | 86.64 | 84.41 | 79.68 | 73.05 | 47.06 | 29.21 |
| M | 93.07 | 92.22 | 76.58 | 53.62 | 86.67 | 85.38 | 73.15 | 70.73 | 42.11 | 19.64 |
| N | 91.99 | 91.68 | 75.14 | 70.18 | 82.52 | 82.59 | 66.30 | 69.54 | 31.90 | 23.40 |
| P | 92.50 | 100 | 64.71 | 75.00 | 100 | 20.00 | 75.00 | 100 | 28.57 | 0 |
| R | 91.93 | 95.80 | 77.42 | 68.27 | 84.87 | 81.13 | 80.25 | 67.74 | 46.48 | 32.00 |
| S | 90.34 | 91.84 | 57.52 | 42.86 | 80.00 | 77.27 | 74.16 | 69.46 | 45.90 | 39.13 |
| ATC class | Share 2010-2013 |  |  |  |  | Relative contribution |  |  |  |  |
| | Pr | $P_I$ | $P_{II}$ | $P_{III}$ | Reg | Pr | $P_I$ | $P_{II}$ | $P_{III}$ | Reg |
| A | 8.12 | 10.06 | 10.67 | 15.44 | 13.91 | 3.29 | 14.35 | 6.57 | 18.40 | 21.74 |
| B | 2.40 | 2.25 | 2.76 | 4.36 | 2.04 | 8.25 | 1.20 | 18.28 | - | - |
| C | 4.24 | 4.04 | 5.47 | 7.02 | 5.01 | 6.35 | 3.41 | 11.67 | - | 9.02 |
| D | 3.75 | 3.64 | 5.91 | 6.13 | 7.05 | 4.45 | 5.96 | 4.17 | 6.07 | 3.36 |
| G | 2.04 | 2.75 | 3.85 | 4.14 | 7.05 | 0.45 | 1.48 | 5.02 | 0.39 | 8.28 |
| H | 0.28 | 0.56 | 0.91 | 1.03 | 1.48 | 2.80 | 0.56 | 4.17 | - | 0.46 |
| J | 13.07 | 8.77 | 7.44 | 9.31 | 9.83 | 25.62 | 5.70 | 18.54 | 15.25 | 1.24 |
| L | 38.08 | 45.88 | 34.90 | 22.75 | 16.51 | 30.12 | 42.18 | 19.41 | 35.13 | 20.88 |
| M | 4.26 | 4.57 | 6.23 | 6.06 | 10.39 | - | 11.41 | 1.98 | 3.87 | 16.68 |
| N | 15.65 | 10.99 | 12.50 | 14.55 | 17.44 | - | 5.91 | - | - | 10.65 |
| P | 0.71 | 0.26 | 0.15 | 0.30 | 0.37 | 4.00 | - | 2.89 | - | 0.76 |
| R | 3.65 | 3.44 | 4.68 | 4.58 | 4.64 | 10.52 | 3.41 | 4.29 | 15.28 | 4.84 |
| S | 3.75 | 2.78 | 4.53 | 4.36 | 4.27 | 4.16 | 4.41 | 3.02 | 5.61 | 2.08 |

**Table S2:** Top-10 disease types by percentage of ongoing clinical studies involving that disease type (source: clinicaltrials.gov)

| Disease type | Percentage of ongoing clinical studies |
| --- | --- |
| Neoplasms | 41.59% |
| Nervous System | 20.96% |
| Cardiovascular | 20.69% |
| Digestive System | 17.29% |
| Immune System | 13.29% |
| Female Urogenital | 13.17% |
| Respiratory Tract | 13.03% |
| Male Urogenital | 10.77% |
| Hemic and Lymphatic | 10.37% |
| Skin and Connective Tissue | 10.36% |

Statistics computed using 33,269 studies whose overall status is “Recruiting” “Active” “Not yet recruiting”, “Enrolling by invitation” or “Available”. We have excluded the studies with only behavioral interventions and the disease type “Pathological conditions, signs and symptoms”. The assignment of a study to a disease type is not exclusive, therefore the reported percentages do not sum to 100.

**Figure S1:**
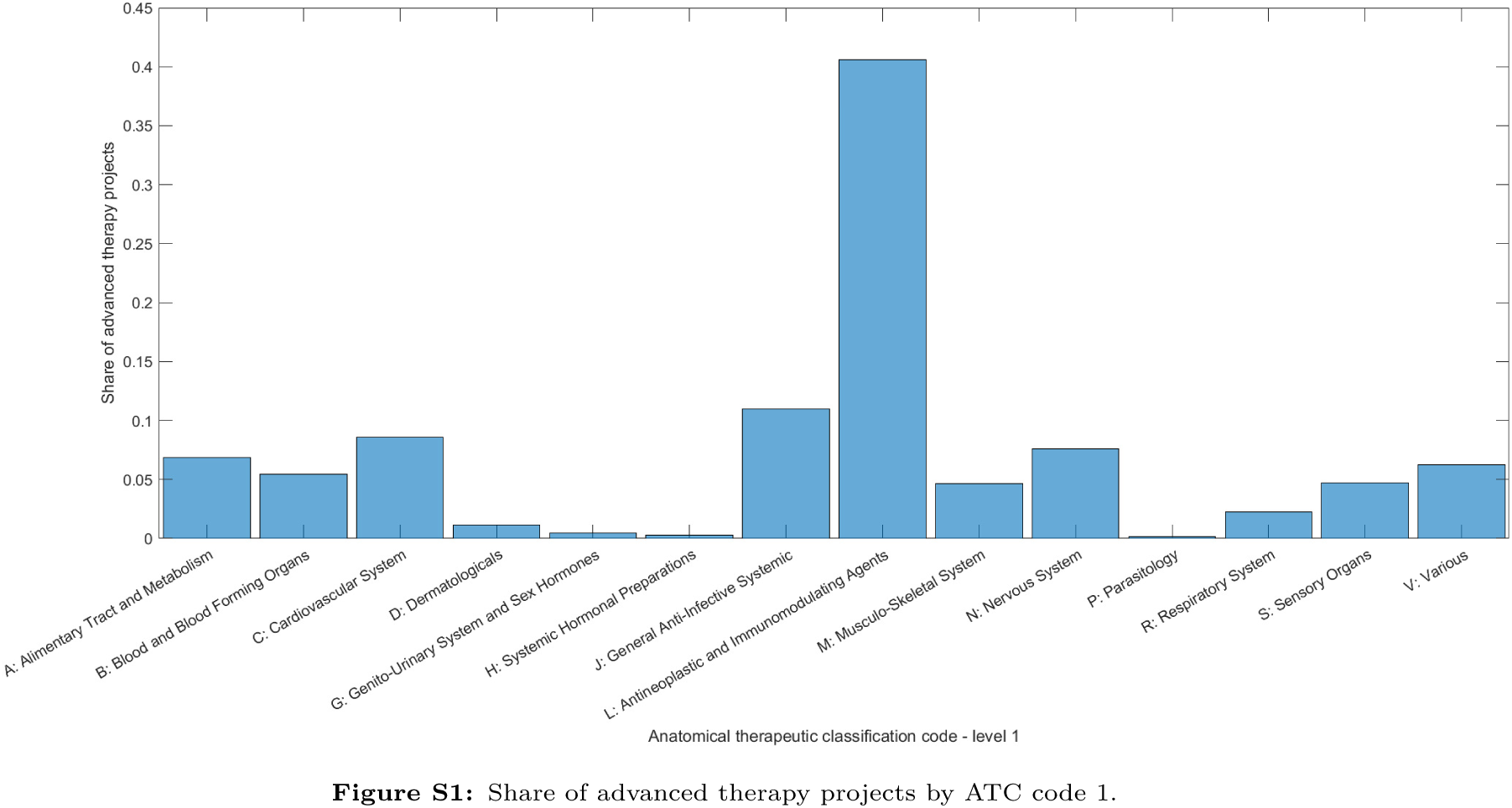
Share of advanced therapy projects by ATC code 1.

**Figure S2:**
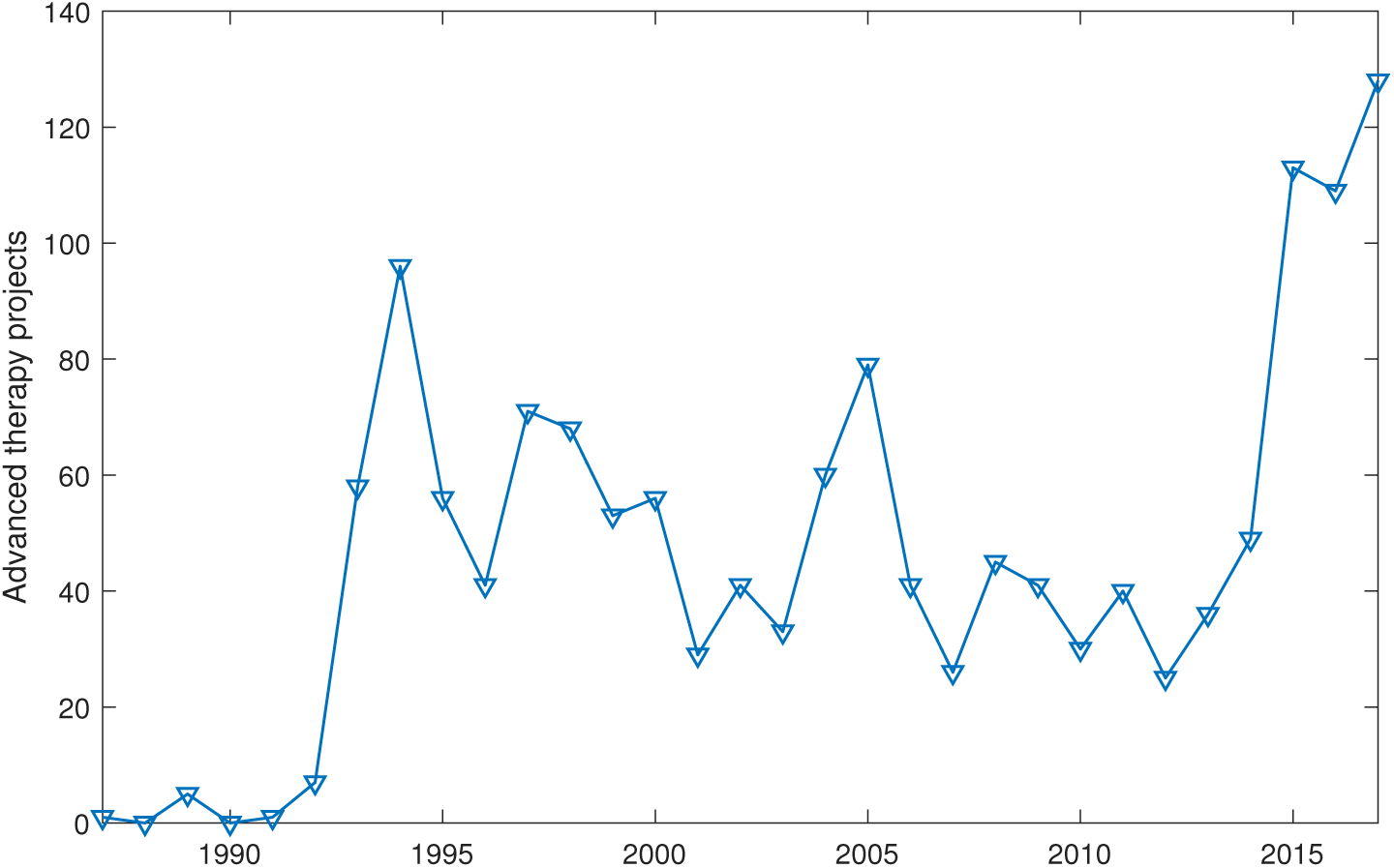
Number of advanced therapy projects by starting year.

**Table S3:** Average (± standard deviation) yearly phase-by-phase attrition rates in three different time intervals (1990-1999, 2000-2009, 2010-2013), for projects focused on treatment of rare diseases. Rare diseases are identified according to a manual classification performed by a domain expert.

| Period | Preclinical | Phase I | Phase II | Phase III | Registration |
| --- | --- | --- | --- | --- | --- |
| 1990-1999 | 87.77(±3.59)% | 52.64(±9.60)% | 76.26(±10.83)% | 61.01(±16.01)% | 46.68(±15.30)% |
| 2000-2009 | 90.83(±2.45)% | 63.91(±7.26)% | 83.15(±2.87)% | 65.64(±13.13)% | 42.04(±14.00)% |
| 2010-2013 | 94.04(±2.02)% | 51.10(±5.23)% | 80.82(±1.78)% | 68.91(±6.01)% | 26.70(±13.21)% |

**Table S4:**
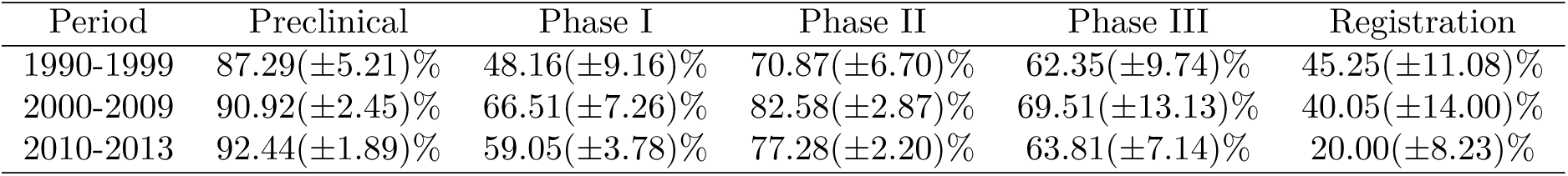
Average (± standard deviation) yearly phase-by-phase attrition rates in three different time intervals (1990-1999, 2000-2009, 2010-2013), for projects for which we have a classification of the institution of the developer (according to the “pharmaceutical”, “biotech” and “non industrial” classification).

| Period | Preclinical | Phase I | Phase II | Phase III | Registration |
| --- | --- | --- | --- | --- | --- |
| 1990-1999 | 87.29($\pm 5.21$)% | 48.16($\pm 9.16$)% | 70.87($\pm 6.70$)% | 62.35($\pm 9.74$)% | 45.25($\pm 11.08$)% |
| 2000-2009 | 90.92($\pm 2.45$)% | 66.51($\pm 7.26$)% | 82.58($\pm 2.87$)% | 69.51($\pm 13.13$)% | 40.05($\pm 14.00$)% |
| 2010-2013 | 92.44($\pm 1.89$)% | 59.05($\pm 3.78$)% | 77.28($\pm 2.20$)% | 63.81($\pm 7.14$)% | 20.00($\pm 8.23$)% |

**Table S5:** Number of projects (percentage share over the whole period) for each OD relationship in the two observation periods (1990-1999 and 2000-2013). *ni*: non-industrial; *ph*: pharmaceutical; *bt*: biotech.

| O \ D | ph | ni | bt |
| --- | --- | --- | --- |
| ph | 772 (40.3%) | - | - |
| ni | 56 (0.3%) | 400 (20.9%) | 102 (5.3%) |
| bt | 135 (7.1%) | - | 451 (23.5%) |

| O \ D | ph | ni | bt |
| --- | --- | --- | --- |
| ph | 1132 (37.1%) | - | - |
| ni | 59 (1.9%) | 510 (16.7%) | 89 (2.9%) |
| bt | 330 (10.8%) | - | 932 (30.5%) |

**Table S6:**
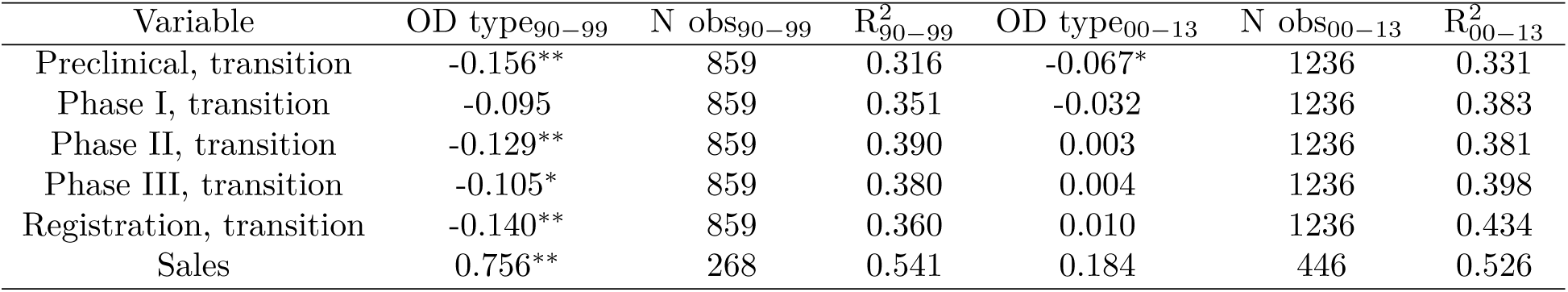
Effect of non-industrial originators vs pharmaceutical developers on project progress and sales.

**Table S7:** Effect of biotech originators vs pharmaceutical developers on project progress and sales.

| Variable | OD type<sub>90-99</sub> | N obs<sub>90-99</sub> | R<sup>2</sup><sub>90-99</sub> | OD type<sub>00-13</sub> | N obs<sub>00-13</sub> | R<sup>2</sup><sub>00-13</sub> |
| --- | --- | --- | --- | --- | --- | --- |
| Preclinical, transition | -0.079 | 925 | 0.287 | -0.039* | 1483 | 0.316 |
| Phase I, transition | -0.011 | 925 | 0.339 | -0.048* | 1483 | 0.359 |
| Phase II, transition | -0.005 | 925 | 0.368 | -0.033 | 1483 | 0.373 |
| Phase III, transition | -0.087* | 925 | 0.371 | 0.013 | 1483 | 0.391 |
| Registration, transition | -0.067 | 925 | 0.359 | 0.042** | 1483 | 0.417 |
| Sales | 0.395 | 269 | 0.505 | -0.355** | 504 | 0.507 |

**Table S8:** Effect of non-industrial originators vs non-industrial developers on project progress and sales.

| Variable | OD type<sub>90-99</sub> | N obs<sub>90-99</sub> | R<sup>2</sup><sub>90-99</sub> | OD type<sub>00-13</sub> | N obs<sub>00-13</sub> | R<sup>2</sup><sub>00-13</sub> |
| --- | --- | --- | --- | --- | --- | --- |
| Preclinical, transition | -0.326** | 1073 | 0.379 | -0.325** | 1578 | 0.449 |
| Phase I, transition | -0.246** | 1073 | 0.403 | -0.264** | 1578 | 0.439 |
| Phase II, transition | -0.217** | 1073 | 0.421 | -0.191** | 1578 | 0.400 |
| Phase III, transition | -0.168** | 1073 | 0.407 | -0.111** | 1578 | 0.393 |
| Registration, transition | -0.162** | 1073 | 0.374 | -0.053** | 1578 | 0.422 |
| Sales | 0.295 | 275 | 0.576 | -0.942** | 456 | 0.572 |

**Table S9:** Effect of non-industrial originators vs biotech developers on project progress and sales.

| Variable | OD type<sub>90-99</sub> | N obs<sub>90-99</sub> | R<sup>2</sup><sub>90-99</sub> | OD type<sub>00-13</sub> | N obs<sub>00-13</sub> | R<sup>2</sup><sub>00-13</sub> |
| --- | --- | --- | --- | --- | --- | --- |
| Preclinical, transition | -0.319** | 1017 | 0.358 | -0.070** | 1334 | 0.336 |
| Phase I, transition | -0.227** | 1017 | 0.384 | -0.088** | 1334 | 0.386 |
| Phase II, transition | -0.223** | 1017 | 0.416 | -0.136** | 1334 | 0.398 |
| Phase III, transition | -0.174** | 1017 | 0.388 | -0.095** | 1334 | 0.406 |
| Registration, transition | -0.175** | 1017 | 0.349 | -0.035 | 1334 | 0.439 |
| Sales | 0.137 | 275 | 515 | -0.581** | 448 | 0.551 |

**Table S10:** Effect of biotech originators vs biotech developers on project progress and sales.

| Variable | OD type<sub>90-99</sub> | N obs<sub>90-99</sub> | R<sup>2</sup><sub>90-99</sub> | OD type<sub>00-13</sub> | N obs<sub>00-13</sub> | R<sup>2</sup><sub>00-13</sub> |
| --- | --- | --- | --- | --- | --- | --- |
| Preclinical, transition | -0.253** | 1299 | 0.326 | -0.056** | 2124 | 0.332 |
| Phase I, transition | -0.206** | 1299 | 0.345 | -0.106** | 2124 | 0.346 |
| Phase II, transition | -0.228** | 1299 | 0.384 | -0.090** | 2124 | 0.303 |
| Phase III, transition | -0.191** | 1299 | 0.363 | -0.080** | 2124 | 0.318 |
| Registration, transition | -0.155** | 1299 | 0.329 | -0.048** | 2124 | 0.337 |
| Sales | 0.362 | 298 | 0.548 | -0.282** | 604 | 0.483 |

## Acknowledgements

We thank Prof. Laura Magazzini for insightful suggestions and technical support.

## Footnotes

1 ClinicalTrials.gov^15^ is a database of privately and publicly funded clinical studies conducted around the world.

## References

1 Pammolli, F., Magazzini, L. & Riccaboni, M. The productivity crisis in pharmaceutical R&D. Nat. Rev. Drug Discov. 10, 428–438 (2011).

2 Scannell, J., Blanckley, A., Boldon, H. & Warrington, W. Diagnosing the decline in pharmaceutical R&D efficiency. Nat. Rev. Drug Discov. 11, 191–200 (2012).

3 Schuhmacher, A., Gassman, O. & Hinder, M. Changing R&D models in research-based pharmaceutical companies. J. Transl. Med. 14, 1–11 (2016).

4 Knight-Schrijver, V., Chelliah, V., Cucurull-Sanchez, L. & Le-Novere, N. The promises of quantitative systems pharmacology modelling for drug development. Comput. Struct. Biotechnol. J. 14, 363–370 (2016).

5 EvaluatePharma. World preview 2018, outlook to 2024. (2018).

6 Garnier, J. Rebuilding the *r*&*d* engine in big pharma. Harv. Bus. Rev. 86, 68–70 (2008).

7 Peck, R. Driving earlier clinical attrition: if you want to find the needle, burn down the haystack. considerations for biomarker development. Nat. Rev. Drug Discov. 12, 289–294 (2007).

8 IQVIA Institute for Human Data Science. Global oncology trends 2018 - innovation, expansion and disruption (2018).

9 IQVIA Institute for Human Data Science. The Changing Landscape of Research and Development (2019).

10 Shapiro, C. Cancer survivorship. N. Engl. J. Med. 2438–2450 (2018).

11 Deloitte. 2018 global life sciences outlook-innovating life sciences in the fourth industrial revolution: Embrace, build, grow (2018).

12 Kepplinger, E. Fda’s expedited approval mechanisms for new drug products. Biotechnol. Law Rep. 34, 15–37 (2015).

13 Arora, A. & Gambardella, A. The changing technology of technological change: general and abstract knowledge and the division of innovative labour. Res. Policy 23, 523–532 (1994).

14 Killick, R., Fearnhead, P. & Eckley, I. A. Optimal detection of changepoints with a linear computational cost. J. Am. Stat. Ass. 107, 1590–1598 (2012).

15 United States National Library of Medicine. Clinicaltrials.gov. http://clinicaltrials.gov. Accessed: 2019-03-14.

16 Aso, E. & Ferrer, I. Cannabinoids for treatment of alzheimer’s disease: moving toward the clinic. Front. Pharmacol. 5, 1–11 (2014).

17 Franco, F. & Cedazo-Minguez, A. Successful therapies for Alzheimer’s disease: why so many in animal models and none in humans? Front. Pharmacol. 5, 1–13 (2014).

18 Miller, K. & Lanthier, M. Investigating the landscape of us orphan product approvals. Orphanet J. Rare Dis. 13, 1–8 (2018).

19 Barham, L. Are the right drugs getting faster FDA approval? (2018). URL http://www.pharmexec.com/are-right-drugs-getting-faster-fda-approval-0.

20 Tufts Center for the Study of Drug Development. Tufts CSDD impact report, may/june 2018 (2018).

21 Jain, E., Kumar, A. Upstream processes in antibody production: Evaluation of critical parameters. Biotechnol. Adv. 26, 46–72 (2018).

22 Stahl, S. Multifunctional drugs: A novel concept for psychopharmacology. CNS Spectrums 14, 71–73 (2009).

23 Van der Schyf, C. The use of multi-target drugs in the treatment of neurodegenerative diseases. Expert Rev. Clin. Pharmacol. 4, 293–298 (2011).

24 Wilcoxon, F. Individual comparisons by ranking methods. Biometrics 1, 80–83 (1945).

25 Barabasi, A.-L., Gulbahce, N. & LoScalzo, J. Network Medicine: A Network-based Approach to Human Disease Nat. Rev. Genet. 12, 56–68 (2011).

26 Arora, A., Gambardella, A., Pammolli, F. & Magazzini, L. A breath of fresh air? firm type, scale, scope, and selection effects in drug development. Manage. Sci. 55, 1638–1653 (2009).

27 Orsenigo, P. F. L. & Riccaboni, M. Technological change and network dynamics. lessons from the pharmaceutical industry. Res. Policy 30, 485–508 (2001).

28 Kirkegaard, H. & Valentin, F. Academic drug discovery centres: the economic and organisational sustainability of an emerging model. Drug Discov. Today 19, 1699–1710 (2014).

29 Magazzini, L., Pammolli, F. & Riccaboni, M. Learning from failures or failing to learn? lessons from pharmaceutical R&D. Eur. Manag. Rev. 9, 45–58 (2012).

30 Belderbos, R., Gilsing, V. & Suzuki, S. Direct and mediated ties to universities: “scientific” absorptive capacity and innovation performance of industrial firms. Strateg. Organ. 14, 32–52 (2016).

31 Hwang, T., Darrow, J. & Kesselheim, A. The FDA’s expedited programs and clinical development times for novel therapeutics, 2012-2016. JAMA 318, 2137–2138 (2017).

32 Nelson, M.R., Tipney, H. & Painter, J.L., Shen, J., Nicoletti, P., Shen, Y., Floratos, A., Sham, P.C., Li, M.J., Wang, J., Cardon, L.R., Whittaker, J.C., Sanseau, P. The support of human genetic evidence for approved drug indications. Nature Genet. 47, 856–860 (2015).

33 Thomsen, S.K., Gloyn, A.L. Human genetics as a model for target validation: finding new therapies for diabetes. Diabetologia 60, 960–970 (2017).

34 Osorio-Mendez, J.F., Cevallos, A.M. Discovery and Genetic Validation of Chemotherapeutic Targets for Chagas’ Disease. Front. Cell. Infect. Microbiol. 8, 439 (2017).

35 Schilsky, R.L., Doroshow, J.H., LeBlanc, M., Conley, B.A. Development and Use of Integral Assays in Clinical Trials. Clin. Cancer Res. 18, 1540–1546 (2012).

36 Hingorani, A.D., van der Windt, D.A., Riley, R.D., Abrams, K., Moons, K.G.M., Steyerberg, E.W., van der Windt, D.A., Schroter, S., Sauerbrei, W., Altman, D.G., Hemingway, H. Prognosis research strategy (PROGRESS) 4: Stratified medicine research. BMJ 346, e5793 (2013).

37 Cassiman, B. & Gambardella, A. Strategic organization of R&D. Adv. Strateg. Manage. 26, 39–64 (2009).

